# Evaluation of deacetylase inhibition in metaplastic breast carcinoma using multiple derivations of preclinical models of a new patient-derived tumor

**DOI:** 10.1101/860205

**Authors:** Tiffany C. Chang, Margarite D. Matossian, Steven Elliott, Hope E. Burks, Rachel A. Sabol, Deniz A. Ucar, Henri Wathieu, Jovanny Zabaleta, Luis De Valle, Sukhmani Gill, Elizabeth Martin, Adam I. Riker, Lucio Miele, Bruce A. Bunnell, Matthew E. Burow, Bridgette M. Collins-Burow

## Abstract

Metaplastic breast carcinoma (MBC) is a clinically aggressive and rare subtype of breast cancer, with similar features to basal-like breast cancers. Due rapid growth rates and characteristic heterogeneity, MBC is often unresponsive to standard chemotherapies; and novel targeted therapeutic discovery is urgently needed. Histone deacetylase inhibitors (DACi) suppress tumor growth and metastasis through regulation of the epithelial-to-mesenchymal transition axis in various cancers, including basal-like breast cancers.

We utilized a new MBC patient-derived xenograft (PDX) to examine the effect of DACi therapy on MBC. Cell morphology, cell cycle-associated gene expressions, transwell migration, and metastasis were evaluated in patient-derived cells and tumors after treatment with romidepsin and panobinostat. Derivations of our PDX model, including cells, spheres, organoids, explants, and *in vivo* implanted tumors were treated. Finally, we tested the effects of combining DACi with approved chemotherapeutics on relative cell biomass.

DACi significantly suppressed the total number of lung metastasis *in vivo* using our PDX model, suggesting a role for DACi in preventing circulating tumor cells from seeding distal tissue sites. These data were supported by our findings that DACi reduced cell migration, populations, and expression of mesenchymal-associated genes. While DACi treatment did affect cell cycle-regulating genes *in vitro,* tumor growth was not affected compared to controls. Importantly, gene expression results varied depending on the cellular or tumor system used, emphasizing the importance of using multiple derivations of cancer models in preclinical therapeutic discovery research. Furthermore, DACi sensitized and produced a synergistic effect with approved oncology therapeutics on inherently resistant MBC.

This study introduced a role for DACi in suppressing the migratory and mesenchymal phenotype of MBC cells through regulation of the epithelial-mesenchymal transition axis and suppression of the CTC population. Preliminary evidence that DACi treatment in combination with MEK1/2 inhibitors exerts a synergistic effect on MBC cells was also demonstrated.

## Background

Metaplastic breast carcinoma (MBC) is a rare breast cancer subtype, comprising 0.45-1% of all breast cancers. This malignancy is molecularly and histologically heterogeneous, expressing both epithelial and mesenchymal features [1–2]; MBC is commonly classified as the triple negative breast cancer (TNBC) PAM50 subtype due to lack of expression of estrogen or progesterone receptors and amplification of the human epidermal growth factor 2 (HER2/Neu) receptor [3]. Poor clinical outcomes are often associated with this cancer diagnosis: the 5-year survival rates for MBC are 38%-78%, compared to 76-93% for invasive ductal carcinoma [4]. A defining feature of MBC is the rapid tumor growth rate; this aggressive clinical presentation contributes to the lower survival rates of patients with MBC compared to patients afflicted with other intraductal carcinomas [4–5]. When compared to another breast cancer subtype that has limited therapeutic targets and high rates of metastasis and relapse, TNBC patients afflicted with MBC have worse disease-free survival and overall survival [6–7]. Despite these differences, MBCs are therapeutically managed similarly to other invasive breast carcinomas, with surgical resection in combination with radiation and/or chemotherapy [8]. However, MBCs have a worse response to neoadjuvant systemic chemotherapy [9] regimens including doxorubicin and cyclophosphamide, or doxorubicin and 5-fluorouracil, in comparison to the overall response of these regimens to other invasive breast cancer subtypes [4,8,10]. The characteristic heterogeneity within MBC, exemplified by the diverse histologic subtypes of MBC, and the dramatically different response rates to chemotherapy of the subtypes [11], contribute to why this malignancy is difficult to manage therapeutically [2, 12]. These findings emphasize the urgent necessity to identify novel therapeutic strategies that are specifically designed to target the unique and heterogeneous nature of MBC.

The implementation of targeted therapies has emerged as a novel approach to treat MBCs, since most MBC cases do not respond to standard systemic chemotherapy regimens. Emerging studies aim to identify novel therapeutic targets for MBC. The MBC transcriptome is similar to the claudin-low spectrum of basal-like breast cancers [13], and next-generation sequencing of 20 MBC tumors that represented various histologic subtypes identified frequent, targetable abnormalities and candidate targets to pursue in MBC including *TP53, PIK3CA, MYC, MLL2, PTEN, CDN2A/B, CCND3, CCNE1, EGFR* and *KDM6A* [14]. A separate study using next-generation sequencing of MBC further supports these findings, as they also demonstrated genetic alterations in the PIK3CA, PTEN and AKT1 mutations were identified in 50% of MBC tumors (18 patients total) and TP53 mutations were found in 56% of tumors [15]. Another group found *PIK3CA* mutations in approximately 50% of MBC tumors (28 patients) [17], and a more recent study supported prevalence of PIK3CA mutations in MBC (present in 23% of MBC tumors; N=297) [18]. Additional targets are being pursued: 14/20 MBC patients had EGFR positive tumors [16], and a high prevalence (39 out of 40) MBC tumors harboring ribosomal protein L39 mutations were found to be susceptible to nitric oxide synthase inhibitors, implicating this as a novel therapeutic strategy for some MBC tumors [19].

Early phase clinical trials of targeted therapies in combination with standard chemotherapy regimens support a role for combination therapy in MBC management. The role of the *PIK3CA/Akt/mTOR* axis in MBC has been demonstrated and it was determined temsirolimus (an mTOR inhibitor) in combination with doxorubicin and bevacizumab resulted in an improved response of MBCs, including complete response [20]. Another example of the potential of combination therapy in MBC treatment is demonstrated by a case study in which a patient with metastatic MBC had a remarkable response to anti-programmed death-ligand 1 (PD-L1) therapy in combination with nab-paclitaxel [21]. Comprehensive profiling of metaplastic breast carcinomas (N=297 samples) revealed a high frequency of PD-L1 overexpression [18]. Furthermore, although MBC is often compared to TNBC subtypes, MBC has distinct therapeutic responses. This is exemplified in a study that demonstrated poor MBC response rate to poly (ADP-ribose) polymerase inhibitor therapy, a targeted therapy which has promising effects in TNBC treatment [22]. A consistent limitation with clinical trials in MBC is that due to the rarity of this malignancy, patient recruitment for larger scale studies and MBC representation in breast cancer research is lacking [10]. Together, these studies show the variability in MBC responses to both targeted and combination treatment and emphasize the importance of establishing more translational MBC models to examine drug effects on this breast cancer subtype.

In this study, we evaluated the potential therapeutic efficacy of histone deacetylase inhibitors (DACi) in MBC. Histone deacetylase enzymes mediate chromatin remodeling, leading to silencing of genes that classically function to suppress tumor growth, inhibit cell-cycle progression, and induce apoptosis in cancer [23]. Paradoxically, this silencing mechanism of action drives tumorigenesis and metastasis. DACi are categorized based on distinct pharmacologic structures: romidepsin (FK228) is a cyclic peptide natural product and a selective HDAC1 and HDAC2 inhibitor, while panobinostat (LBH589), a nonselective deacetylase inhibitor, is a cinnamic hydroxamic acid analog of M-carboxycinnamic acid bishydroxamate [24]. These DACi have been investigated as targeted therapies for select cancer types: romidepsin and vorinistat are approved to treat cutaneous T-cell lymphoma [25], belinostat is approved to treat peripheral T-cell lymphoma [26], and panobinostat is approved to treat multiple myeloma [27]. Additionally, DACi therapies are in various stages of clinical trials for other cancer types, either as single agent or in combination therapies [28]. Examples include the combination of the DACi abexinostat with pazopanib in advanced renal cell carcinoma [29], the DACi vorinistat in combination with bevacizumab in clear-cell renal cell carcinoma [30], and the DACi abexinostat in combination with doxorubicin in metastatic sarcoma [31]. Other tumor types in which DACi therapy is being investigated include lung, pancreatic, advanced colorectal and hepatocellular carcinoma, Hodgkin lymphoma, hematologic malignancies, and breast cancer [28].

DACi have emerged as an effective targeted therapy in breast carcinomas in preclinical studies [32], specifically in TNBC subtypes [33]. Our laboratory has extensively evaluated DACi in TNBC cells and tumors [34–36]; we observed different biologic effects on tumorigenesis and metastasis with pan-DAC inhibition compared to class-specific inhibitors [35]. We have previously shown that aggressive breast cancer subtypes, such as TNBCs [34–36], specifically claudin-low TNBCs [36], are susceptible to DACi. Treatment with romidepsin and panobinostat reversed the mesenchymal phenotype of TNBCs, in addition to suppressing tumorigenesis and metastasis [35], consistent with findings from other groups [33, 37]. Maintenance of a mesenchymal phenotype is important in TNBC biology; the process through which cells acquire these cellular characteristics, also known as the epithelial-to-mesenchymal transition (EMT), is one proposed mechanism that is integral in the initiation of metastasis. In EMT, luminal breast cancer cells lose epithelial characteristics (cytoskeletal rearrangement, loss of epithelial markers) and gain mesenchymal features (invasiveness, formation of cellular protrusions, expression of mesenchymal markers). This alteration facilitates detachment of the cells from the primary tumor and intravasation into the vasculature, through which the cells migrate and seed distal tissue sites [36]. Some studies have described the acquisition of mesenchymal characteristics contributing to tumor-initiating capabilities of cancer cells and tumor development [39].

The associations amongst EMT, metastasis, and chemotherapeutic resistance are not completely understood. Expression of EMT-associated genes was initially thought to drive epithelial stem cell properties and breast cancer metastasis [41–43]. Other hypotheses suggest that EMT is not required for metastasis [44] and that epithelial- and mesenchymal-like states are mediated by the tumor microenvironment [45]. Despite this lack of congruency, targeted therapies that reverse the mesenchymal phenotype of cancer cells have been shown to improve the sensitivity of tumors to cytotoxic effects of chemotherapeutic agents [44,46,47]. Our group has previously demonstrated a similar role for DACi in combination therapy to re-sensitize endocrine-resistant breast tumors to therapeutic agents [40]. DACi treatment sensitizes TNBC cells to both targeted and systemic chemotherapies, especially DNA-damaging agents [17,48,49]. For example, DACi induced BRCA1 expression in TNBC cells and exhibited synergistic lethality with poly-ADP-ribose polymerase (PARP) inhibitor and cisplatin [50]. Another study found the DACi suberoylanilide hydroxamic acid to enhance anti-tumor effects of the PARP inhibitor olaparib in TNBC cells [51]. In this manuscript we aim to evaluate the therapeutic efficacy of DACi on the mesenchymal and metastatic phenotype of MBC and determine synergistic cytotoxic effects of DACi on MBC cells in combination with other targeted therapeutics.

Patient-derived xenograft (PDX) models are more translational than immortalized cell line-derived models because they maintain features that are present in the original patient tumor [52, 53]. Because MBC is a rare malignancy, there are fewer available model systems to study this disease compared to other breast cancer subtypes; it is crucial to develop new PDX models for MBC. Our group has established and characterized a new PDX MBC model, TU-BcX-4IC. Evaluating therapeutic response on various aspects of a patient’s tumor is crucial to comprehensively understanding the mechanism of action on the complex biological systems that comprise MBC tumors: the cell population, cell clusters in suspension, cells in interaction with the extracellular matrix both *in vitro* (using organoids) and *ex vivo* (using tumor explants), and the intact tumor *in vivo.* Emerging studies show different responses of targeted drugs on cell viability and gene expression in adherent conditions compared to cells plated in suspension conditions [54, 55] and *ex vivo* treatments [56]. We hypothesize that by evaluating drug effects more comprehensively in the laboratory setting using this systems-based approach, there will be an improvement in the translation of preclinical observations into clinical practice.

There are currently no preclinical nor clinical studies that evaluate the role of DACi in MBC, to our knowledge. In this study we demonstrate the effect of histone deacetylase inhibitors on the mesenchymal phenotype and metastatic potential in a new PDX model derived from a clinically aggressive and drug-resistant MBC.

## Materials and Methods

### Reagents and cell lines utilized

Human TNBC cell lines were obtained from American Type Culture Collection (ATCC). TU-BcX-4IC cells were established from the original TU-BcX-4IC tumor explant, before implantation into mice for propagation. TU-BcX-4IC was dissected and a small tumor explant was plated in a 10cm dish in 10% DMEM media (described below). TU-BcX-4IC cells grew out from the tumor and were propagated until a cell line was established. Cells were grown in DMEM supplemented with 10% Fetal bovine serum (FBS), insulin, non-essential amino acids, minimal essential amino acids, antibiotic-antimycotic, and sodium pyruvate at 37°C in 5% CO_2_. Dulbecco’s modified Eagle’s medium and non-essential amino acids were purchased from Caisson Laboratories (Smithfield, UT). Minimum essential amino acids, sodium pyruvate, antibiotic-antimycotic, insulin, TrypLE were purchased from Invitrogen (Carlsbad, CA). Dimethyl sulfoxide was purchased from Thermo Fischer Scientific. FBS was obtained from Gemini Bio-Products (West Sacramento, CA). Romidepsin was purchased from ApexBio (Houston, TX). Panobinostat was generously provided by Novartis Pharmaceutical Inc. (East Hanover, NJ).

### Generation of cell line-derived spheres

MDA-MB-231 and TU-BcX-4IC cells were plated in suspension conditions in low-attachment 96-well plates (1,000 cells per well), with serum-free F12/DMEM (Sigma-Aldrich, Carlsbad, CA; Cat. No D6421) media supplemented with B-27 (ThermoFisher Scientific, Plaquemine, LA; Cat. No. 17504044). Cells were maintained in cell culture conditions for 24 hours to facilitate sphere formation. After 24 hours, suspended cells/spheres were treated with drug and DMSO controls for one week, with treatment frequencies of every three days. Area of spheres were used one endpoint to quantify drug effects on sphere formation. For these quantifications, images were obtained of spheres at both 40X and 100X magnifications, depending on the average size of the spheres. The lengths and widths of individual spheres after treatments were measured using the Aperio ImageScope program. The area was calculated and recorded and compared to DMSO treatment controls.

### Crystal violet staining and dose/response quantification

TU-BcX-4IC primary cell line established from the 4IC PDX tumor model, and MDA-MB-231 cells were utilized. Cells were maintained in DMEM supplemented with 10% FBS, insulin, non-essential amino acids, minimal essential amino acids, antibiotic-antimycotic, and sodium pyruvate at 37°C in 5% CO_2_. TU-BcX-4IC cells were exposed to charcoal-stripped media (phenol-free DMEM supplemented with 5% charcoal-stripped FBS (Gibco Invitrogen, Carlsbad CA), non-essential amino acids, minimal essential amino acids, Glutamax (Gibco Invitrogen, Carlsbad CA), penicillin-streptomycin (100 U/mL) for 24 hours. The following day, cells were treated with various concentrations of romidepsin, panobinostat, or DMSO treatment control. After three days, cells were fixed with glutaraldehyde and stained with crystal violet to observe the response to chemotherapies. Representative images were captured with brightfield microscopy. The cells remaining were quantified by lysis with 33% acetic acid and absorbance was measured at 620 nm wavelength with a spectrophotometer.

### Cell migration assays

TU-BcX-4IC cells were exposed to charcoal-stripped media (phenol-free DMEM supplemented with 5% charcoal-stripped FBS, non-essential amino acids, minimal essential amino acids, Glutamax, penicillin-streptomycin) for 24 hours. The following day, cells were treated with romidepsin (100 nM), panobinostat (100 nM), or DMSO treatment control (0.01% DMSO). After 48 hours of treatment, cells were counted using an automatic cell counter and 25,000 cells were plated on top of a migration chamber with 0.8-micron pores in serum-free Opti-MEM media (Gibco, USA). DMEM media (10% FBS) with supplements used for cell culture maintenance was plated in the bottom chamber. Cells were treated in-well for an additional 24 hours. Then, migration membranes were rinsed, scrubbed, fixed with formalin and stained with crystal violet. Membranes were mounted on slides and stained cells fixed to the membranes were quantified using the ImageJ software. Data is represented as mean ± standard error of the mean (SEM).

### Patient-derived xenografts

TU-BcX-4IC was derived from a mastectomy specimen of a 57-years-old female Caucasian patient with a metaplastic breast carcinoma unresponsive to neoadjuvant adriamycin/cyclophosphamide therapy. TU-BcX-4IC was propagated in SCID/Beige mice (CB17.cg-PrkdcscidLystbg/Crl) obtained from Charles River. The tumor specimen was sectioned into 5×5 mm pieces under sterile conditions and coated in Matrigel (Cat No. 354234, Fisher Scientific, Waltham, MA, USA) to promote tumor take. After the first initial passage, Matrigel was no longer used, to show that TU-BcX-4IC grew independently of Matrigel. Mice were given subcutaneous injections of Melocixam (5 mg/kg) prior to surgery. To passage the tumor, TU-BcX-4IC was implanted bilaterally into mammary fat pads of female SCID/Beige mice under anesthesia with a mixture of isoflurane and oxygen. Tumor volume was measured biweekly using digital calipers. Once tumor volume reached 1000 mm^3^, mice were euthanized, and the tumors were harvested for subsequent passage. Lungs and liver were also harvested, formalin-fixed and paraffin-embedded to observe baseline metastases. Mice were euthanized with CO_2_ and cervical dislocation.

This study was carried out in strict accordance with the recommendations in the Guide for the Care and Use of Laboratory Animals of the National Institutes of Health. The protocol was approved by the Tulane University Internal Review Board for Research on Human Tissue, IACUC (Protocol Number: 635). All procedures performed in studies involving SCID/Beige mice were approved by the Tulane University Institutional Animal Care and Use Committee and in accordance with ethical standards. All surgery was performed under isoflurane and oxygen anesthesia, and all efforts were made to minimize suffering.

### Generation of PDX-Os and live/dead fluorescence stain

When SCID/Beige mice implanted with TU-BcX-4IC tumors were passaged to maintain the tumor, small (2×2 mm^2^) tumor pieces were plated in suspension culture conditions. Cells were allowed to grow out from the tumor pieces for 7 days, which contained mixed cell and stromal populations. At that time, PDX-Os were transferred to a 96-well ULA spheroid plate (Corning, NY, Cat. No. 4515) and treated with romidepsin or DMSO control. After 72 hours, media was removed and spheres were stained using the PromoKine live/dead staining kit (New York, USA). Cells were exposed to Calcein-AM (2µM) and Ethidium homodimer-III (5µM) mixed with phosphate buffered saline. Calcein-AM can be transported through the cell membrane of live cells, where fluorescence activation is based on interaction with esterase enzymes. Ethidium homodimer binds to DNA of lysed (dead) cells. Cells were incubated for 45 minutes. Stained cells were imaged with confocal fluorescence microscopy and images were captured (8 images per well of adherent cells, 5 images per well of low-suspension cells). The 588 nM excitation channel was used to identify red, ‘dead’ cells, and the 420 nM excitation channel was used to visualize green, ‘live’ cells. Representative images were taken at 100x magnification.

### Immunofluorescence staining for CSC markers

TU-BcX-4IC cells were plated as described in the crystal violet staining section. Cells treated for 72 hours with DMSO, romidepsin (100nM) or panobinostat (10nM). Treated cells were fixed in formalin (10% buffered formalin phosphate, Fischer Scientific, Hampton NH) and permeabilized with Triton-X100 (MP Biomedicals, St. Ana CA). A primary conjugated antibody against CD44 (BioLegend, Cat No. 103006, San Diego, CA) was utilized to stain the cells, and DAPI (NucBlue Fixed Cell Stain ReadyProbe, Life Technologies, Carlsbad CA) was utilized to highlight nuclei. ApoTome (commercial structure illumination microscopy by Zeiss, Thornwood, NY) fluorescent images were captured on an inverted microscope (Zeiss) and digitally filtered to obtain optical slices. 5 images per well were captured at 200x, n=3.

### RNA sequencing

After validating the integrity of the RNA samples in an Agilent BioAnalyzer 2100, 200 ng total RNA were used to make cDNA libraries using the TruSeq RNA Exome kit following the vendor’s recommendations (Illumina) and sequenced at 2 x 75 bp in a NextSeq500 instrument (Illumina). FASTQ files were aligned to the *Homo sapiens*/hg19 reference genome using the RNA-seq Alignment tool v1.1.1 in the Illumina’s BaseSpace. Raw counts were extracted and used to find genes differentially expressed using DESEQ2 v1.16.1 in R-Studio 1.1.383. The data was normalized using the Variant Stabilizing Transformation and log2. A p value of <0.05 was considered significant and was further corrected by FDR < 0.05. Heatmaps were built using the *Pretty Heatmaps* application (pheatmaps) v1.0.10 in R-Studio.

### Quantitative real time polymerase chain reaction

Total RNA was isolated and extracted from 4IC tumor samples and TU-BcX-4IC adherent cells using Quick-RNA MiniPrepTM (Zymo Research, Irvine, CA) according to manufacture protocol. After confirming RNA quality and quantity, RNA was reverse-transcribed (2 ug) into cDNA (iScript kit, BioRad Laboratories, Hercules, CA) and analyzed by qRT-PCR. Cycle number was normalized to β-actin and vehicle-treated cells scaled to 1, n = 3. Primers (Invitrogen, Carlsbad, CA) were generated with sequences as follows: *β-actin* F--5’-GGCACCCAGCACAATGAAGA-3’; *β-actin* R-5’-ACTCCTGCTTGCTGATCCAC-3’; *CDH1* F-5’-AGGTGACAGAGCCTCTGGATAGA-3’, *CDH1* R-3’-TGGATGACACAGCGTGAGAGA-3’; *VIM* F-5’-GAGAACTTTGCCGTTGAAGC-3’, *VIM* R-5’-GCTTCCTGTAGGTGGCAATC-3’; *CDH2* F-5’-GCCCCTCAAGTGTTACCTCAA-3’, *CDH2* R-5’-AGCCGAGTGATGGTCCAATTT-3’; *ZEB1* F-5’-TGCACTGAGTGTGGAAAAGC-3’, *ZEB1* R-5’-TGGTGATGCTGAAAGAGACG-3’; *ZEB2* F-5’-CGCTTGACATCACTGAAGGA-3’, *ZEB2* R-5’-CTTGCCACACTCTGTGCATT-3’; *PLAU* F-5’-GGAAAACCTCATCCTACACAAGGA-3’, *PLAU* R-5’-CGGATCTTCAGCAAGGCAAT-3’; *FOS* F-5’-GAATGCGACCAACCTTGTGC-3’, *FOS* R-5’-AGGGATCAGACAGAGGGTGT-3’; *FRA-1* F-5’-CGAAGGCCTTGTGAACAGAT-3’, *FRA-1 R*-5’-CTGCAGCCCAGATTTCTCA-3’.

### Treatment of cells with National Cancer Institute-approved oncology drug set

For these experiments, TU-BcX-4IC primary cell line established from the 4IC PDX tumor model, and MDA-MB-231 cells were utilized. Cells were maintained in DMEM supplemented with 10% FBS, insulin, non-essential amino acids, minimal essential amino acids, antibiotic-antimycotic, and sodium pyruvate at 37°C in 5% CO_2_. TU-BcX-4IC cells were seeded in 96-well plates and treated with the commercially available NCI oncology drug panel (https://wiki.nci.nih.gov/display/NCIDTPdata/Compound+Sets). After three days, cells were fixed with glutaraldehyde and stained with crystal violet to observe the response to chemotherapies. Representative images were captured with brightfield microscopy. Stained cells were quantified two ways: 1) crystal violet-stained cells were lysed with 33% acetic acid and absorbance was measured at 620 nm wavelength with a spectrophotometer or 2) visible cells remaining after treatment were quantified using the ImageJ program. For the ImageJ quantification, images were captured at 100X using brightfield microscopy, and three images per 96-well plate were quantified for each treatment group. Debris were not included in the quantification.

### Ex vivo treatment of PDX tumor pieces

TU-BcX-4IC was resected from mice after passages 2 and 6 (T2 and T6) and dissected into 5×5 mm^3^ pieces. Tumor pieces were plated in 12-well plates with 10% FBS/DMEM media and treated with romidepsin (100 nM) or DMSO control (0.1%) for 72 hours. TU-BcX-4ICT2 tumors were treated in duplicate due to limited tissue availability and TU-BcX-4ICT6 tumors were treated in triplicate. PDX-Es were collected after 72 hours, and RNA was extracted using Qiazol Lysis Reagent (Cat No. 79306; Qiagen, Valencia, CA, USA) and dissection of the tumor with scissors. Total RNA was isolated and cDNA was made as described above. mRNA was analyzed by qRT-PCR.

### In vivo treatment of TU-BcX-4IC xenografts

Female SCID/Beige mice (n=5/group) were inoculated with TU-BcX-4IC tumor implants (3×3 mm^3^) in the mammary fat pads. For these studies, mice were treated with romidepsin (0.25 mg/kg) or DMSO vehicle control twice a day. This dose was generated from pharmacokinetic and pharmacodynamic data. Treatments were initiated immediately after tumor implantation (after 24 hours) due to the rapid growth rate and tumor take of this PDX model. After tumors reached 850 mm^3^ in volume, mice were euthanized. Lungs and livers were harvested to examine metastasis, and peripheral blood was collected.

### Immunohistochemistry staining

Tumor specimens, lungs, and livers were harvested at necropsy, and fixed in formalin. The samples were sent to our Department of Pathology at Tulane University where they were paraffin-embedded, sectioned and stained with hematoxylin and eosin (H & E). Lungs and liver sections were imaged using the Aperio Scanscope instrument (Aperio Technologies, Inc., Vista, CA, USA). Quantification of metastasis to the lungs and livers was achieved using ImageScope software (Aperio Technologies, Inc.).

### Flow cytometry for cancer stem cell populations

Circulating tumor cells were collected in whole blood with 0.5M EDTA (Gibco Invitrogen, Carlsbad CA), incubated in red blood cell lysis buffer (0.008% NH4Cl, pH 7.2-7.4; Sigma-Aldrich, St. Louis MO), and washed with PBS. Collected cells from the tumor and blood samples were placed in staining solution containing 1% Bovine Serum Albumin (BSA; Sigma-Aldrich) and 1% CD16/CD32 Mouse BD Fc BlockTM (BD Biosciences) in PBS. The following primary antibodies were used: Anti-human CD24 (APC), and anti-human/mouse CD44 (PE-Fluor 610) purchased from eBiosciences (San Diego, CA, USA). All cells from the blood were analyzed with a Galios Flow Cytometer (Beckman Coulter, Brea, CA, USA) running Kaluza software (Beckman Coulter). At least 5000 events were analyzed and reported as the mean ± SEM.

### Hematoxylin & eosin staining

Livers, lungs, and tumor tissues were processed by the Department of Histology and Pathology at Tulane University. As per standard protocol, formalin-fixed tissues were paraffin-embedded, sectioned at 4 µM, and mounted on glass slides. Mounted sections were then exposed to xylene, ethanol, and acetic acid with intermittent washings with water before being stained with hematoxylin and eosin. After staining, slides were then again exposed to ethanol and xylene to complete the protocol.

### Statistical Analysis

The data was analyzed through unpaired Student’s t-tests performed in Prism v7 (Graphpad, Inc.) with p-values of < 0.05 considered statistically significant. Error bars are represented as mean ± standard error of mean (SEM).

## Results

### DAC inhibition suppresses viability of TU-BcX-4IC cells in a dose-dependent manner and reverses mesenchymal morphology

To understand the role of histone DACi in metaplastic breast cancer, we first evaluated drug effects on cell viability and cell morphology of TU-BcX-4IC cells derived from an MBC patient tumor established in our laboratory. Cells were plated in adherent conditions and treated with panobinostat, romidepsin, or DMSO control for 72 hours. Cell viability was determined when crystal violet-stained cells were lysed, and absorbance was quantified. Both panobinostat and romidepsin reduced cell viability in dose-dependent manners, compared to DMSO control treatments (Figs 1A-1C); panobinostat and romidepsin had IC50 values of 0.1 µM and 0.5 µM, respectively (Fig 1B). Alterations in cell morphology were also observed in the DACi-treated cells. DAC inhibition reversed the mesenchymal cell morphology and increased epithelial features; the mesenchymal morphology is characteristic of the more metastatic TNBC subtypes. Suppression of the mesenchymal morphology is defined by fewer cells with protrusions, enhanced cell-cell contact, and rounder cell shapes [46]. Transwell migrations of cells treated for 72 hours with DMSO, romidepsin, and panobinostat showed decreased migration at 100 nM. Romidepsin treatment more significantly decreased cell migration, compared to panobinostat and the DMSO control (Fig 1D).

**Fig 1.**
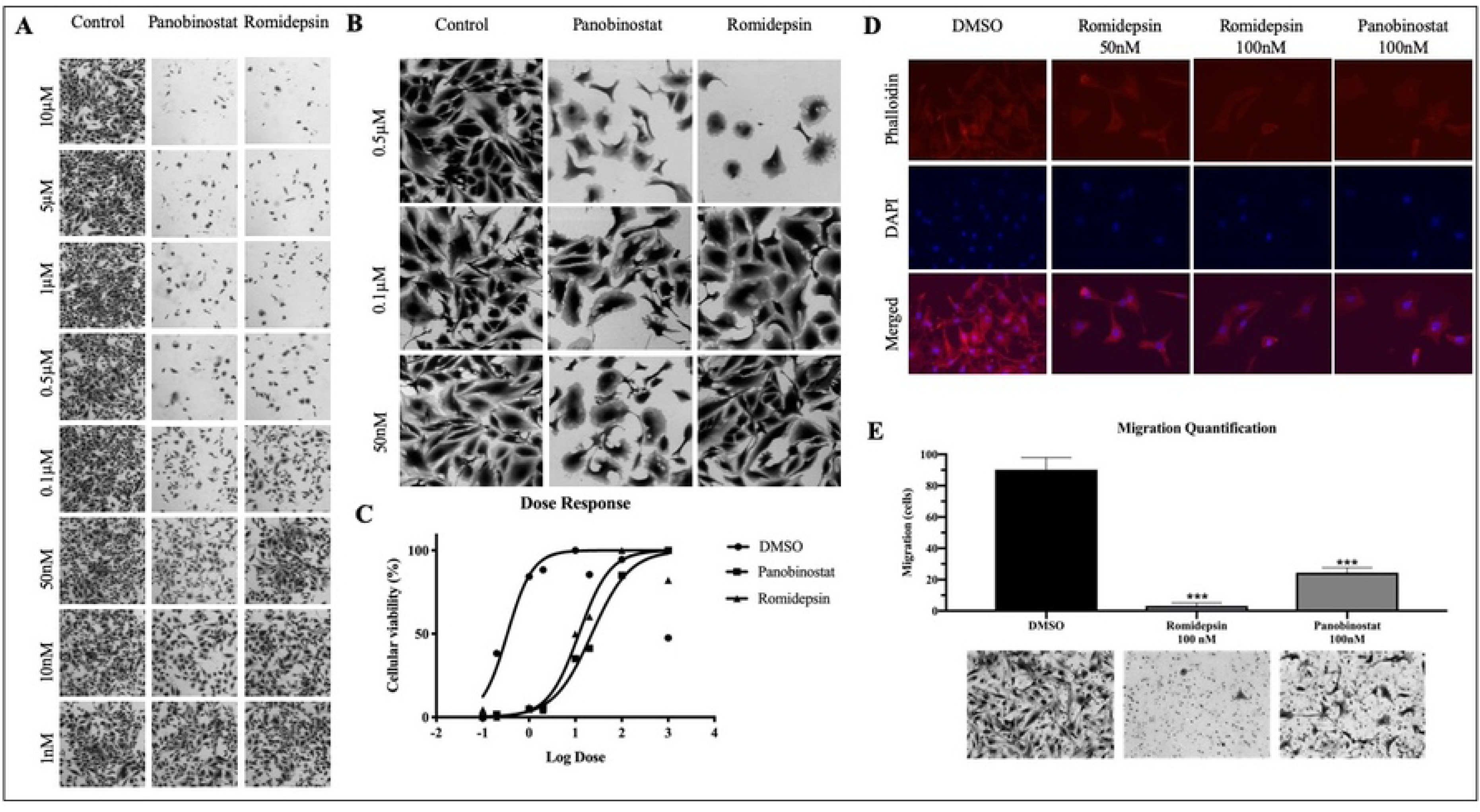
Dose-dependent response to panobinostat or romidepsin in adherent TU-BcX-4IC cells. TU-BcX-4IC cells were plated and treated with serial dilutions of panobinostat, romidepsin, or DMSO control for 72 hours. Cell viability and morphology were compared to DMSO control. Images were captured at 4X and 10X. (A) Panobinostat and romidepsin treatment to TU-BcX-4IC adherent cells demonstrate cytoxicity at 50 µM and 0.1 µM, respectively. Representative images are shown at 4X magnification. (B) Panobinostat and romidepsin treatment reverse the mesenchymal transition and promote epithelial-like characteristics. Representative images are shown at 10X magnification. (C) Dose-response analysis of TU-BcX-4IC treated with panobinostat, romidepsin, or DMSO shows increased potency and efficacy of romidepsin compared to panobinostat. After crystal violet staining, plates were lysed and absorbance was measured (570 nm) and normalized to DMSO controls to quantify drug response. (D) Transwell migration of TU-BcX-4IC cells pre-treated for 72 hours with DMSO, romidepsin or panobinostat (24 hours migration, 100 nM treatments). Error bars are represented as standard error of mean. *** p <0.001.

Patient-derived xenograft organoids (PDX-O) have emerged as a translational model to evaluate cytotoxicity because they recapitulate the microanatomy of patient tumors. These models facilitate testing drug effects on a more translational model, in which cells solely grown in suspension culture. Furthermore, organoids have been shown to more accurately predict clinical response to therapeutics, compared to cell line studies [57, 58]. We examined romidepsin effects in TU-BcX-4IC PDX-Os. PDX-Os were established from TU-BcX-4IC PDX tumor explants and maintained in suspension culture conditions. PDX-Os were treated with romidepsin (100 nM) or DMSO control for 72 hours and stained with a live/dead kit; fluorescence microscopy was used to visualize relative live and dead cell populations. Romidepsin treatment resulted in increased cytotoxicity of cells in the organoids compared to DMSO controls (Fig 2A). This experiment was performed in quadruplicate, and all treated PDX-Os are shown in Fig 2A.

**Fig 2.**
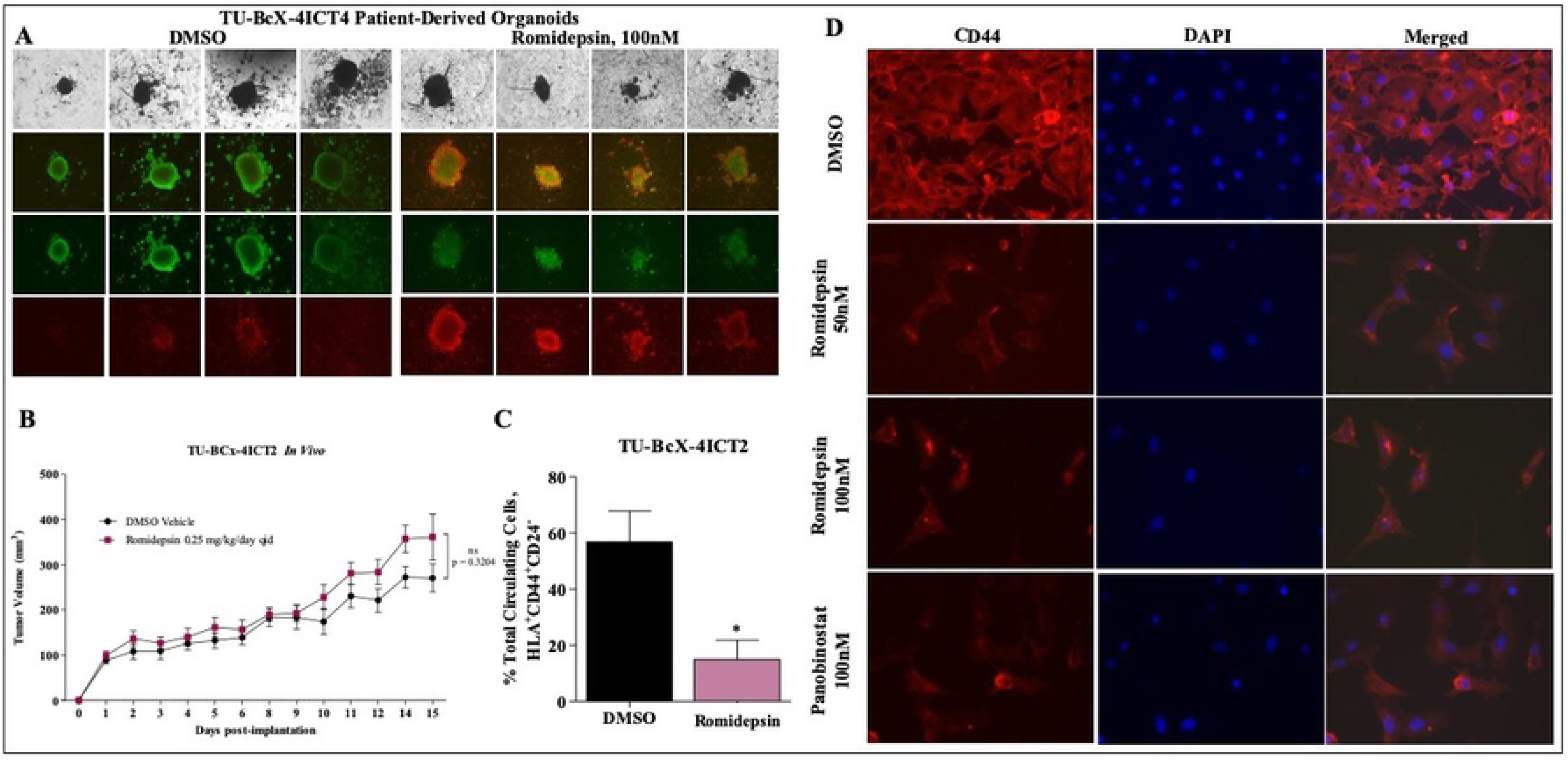
Romidepsin cytotoxicity to TU-BcX-4ICT4 patient-derived organoids and *in vivo* effects on tumorigenesis and circulating tumor cells. (A) PDX-Os were generated from TU-BcX-4ICT4 in 3D culture conditions and treated for 72 hours with romidepsin (100 nM) or DMSO control. PDX-Os were treated with a live (Calcein AM)/dead (EthD II) immunofluorescent staining kit to highlight live (green) or dead (red) cells. Representative images were captured at 40X magnification (brightfield images) or 50X magnification (immunofluorescent images). (B) TU-BcX-4IC tumor pieces (passage 2; T2) were implanted bilaterally in the mammary fat pads of SCID/Beige mice treated with romidepsin or DMSO vehicle. Tumor volume was measured biweekly with calipers and after 20 days, tumors were excised and mice were sacrificed. (C) Twenty days after tumor excision, peripheral blood was collected to evaluate circulating tumor and stem cell populations, defined as HLA^+^CD44^+^CD24^-^. Percentage of circulating tumor cells were significantly reduced in mice treated with romidepsin compared to those treated with DMSO. Error bars represent standard error of mean. *p <0.05.

### DAC inhibition does not affect tumor growth but suppresses circulating tumor cell populations in vivo

To assess the effect of DACi on tumor formation, TU-BcX-4IC PDX explants were implanted in SCID/Beige mice and mice were intraperitoneally treated with romidepsin (0.25 mg/kg/day) or DMSO vehicle. Notably, at baseline, TU-BcX-4IC exhibits rapid tumor growth rates, compared to other TNBC models established by our group. No significant change in tumor growth was observed in romidepsin-treated tumors (194.0 ± 26.86 N=14) compared to DMSO-treated control tumors (160.0 ± 20.07 N=14) (Fig 2B). Tumors were excised, and mice were monitored for an additional 20 days to evaluate metastasis. Histopathologic analysis of excised tumors revealed cancer cells with pleomorphic nuclei, mitotic figures, and hyperchromaticity in both romidepsin and DMSO-treated mice (S1 Fig). After 20 days, mice were euthanized and peripheral blood was harvested to examine the effect of DACi on levels of cancer stem cells within the circulating tumor cell (CTC) populations. CTCs were identified as HLA^+^ (a human-specific marker), and CD44^+^CD24^-^ cells indicate cancer stem cells [59]. We found a significantly reduced percentage of circulating tumor and stem cells in the peripheral blood of romidepsin-treated mice compared to DMSO-treated mice (Fig 2C). While these data show that romidepsin does not affect tumor formation, romidepsin suppresses the cancer stem cell phenotype that drives metastasis. These data were confirmed by our findings that CD44 expression was less robust in TU-BcX-4IC cells after treatment with panobinostat and romidepsin, using immunofluorescence staining (Fig 2D).

### Romidepsin treatment suppresses metastasis in TU-BcX-4IC

Given our findings that romidepsin suppresses CTC populations in peripheral blood, we next evaluated the effect of romidepsin on TU-BcX-4IC metastasis in immunocompromised mice. Notably, at baseline TU-BcX-4IC is a highly metastatic PDX model with lesions to both lungs and livers. Lungs and livers were harvested, formalin-fixed, paraffin-embedded and H & E-stained to visualize and quantify metastatic lesions. Romidepsin significantly reduced the number of metastatic lesions to the lungs (DMSO: 45.75 ± 13.98 N=4, Romidepsin: 10.33 ± 2.028 N=3) (Fig 3A), although the total area of lung metastatic lesions did not change significantly between the treatment and non-treatment groups (DMSO: 38930 ± 33090 N=4, Romidepsin: 14920 ± 13180 N=3) (Fig 3B). Although romidepsin reduced the total number and total area of liver metastases, neither the total number of metastatic lesions (DMSO: 26.00 ± 11.68 N=3, Romidepsin: 15.00 ± 3.000 N=3) (Fig 3D), nor the total area of metastases (DMSO: 887.8 ± 128.1 N=3, Romidepsin: 776.8 ± 88.59 N=3) (Fig 3E) were significantly reduced. Representative images of H & E-stained lungs (Fig 3C) and livers (Fig 3F) are included to demonstrate the size of metastatic lesions in the DMSO and romidepsin treatment groups. Together, these data show that although romidepsin treatment decreased the number of lung metastases of the TU-BcX-4IC PDX model, romidepsin treatment did not reduce overall areas of lung and liver metastasis.

**Fig 3.**
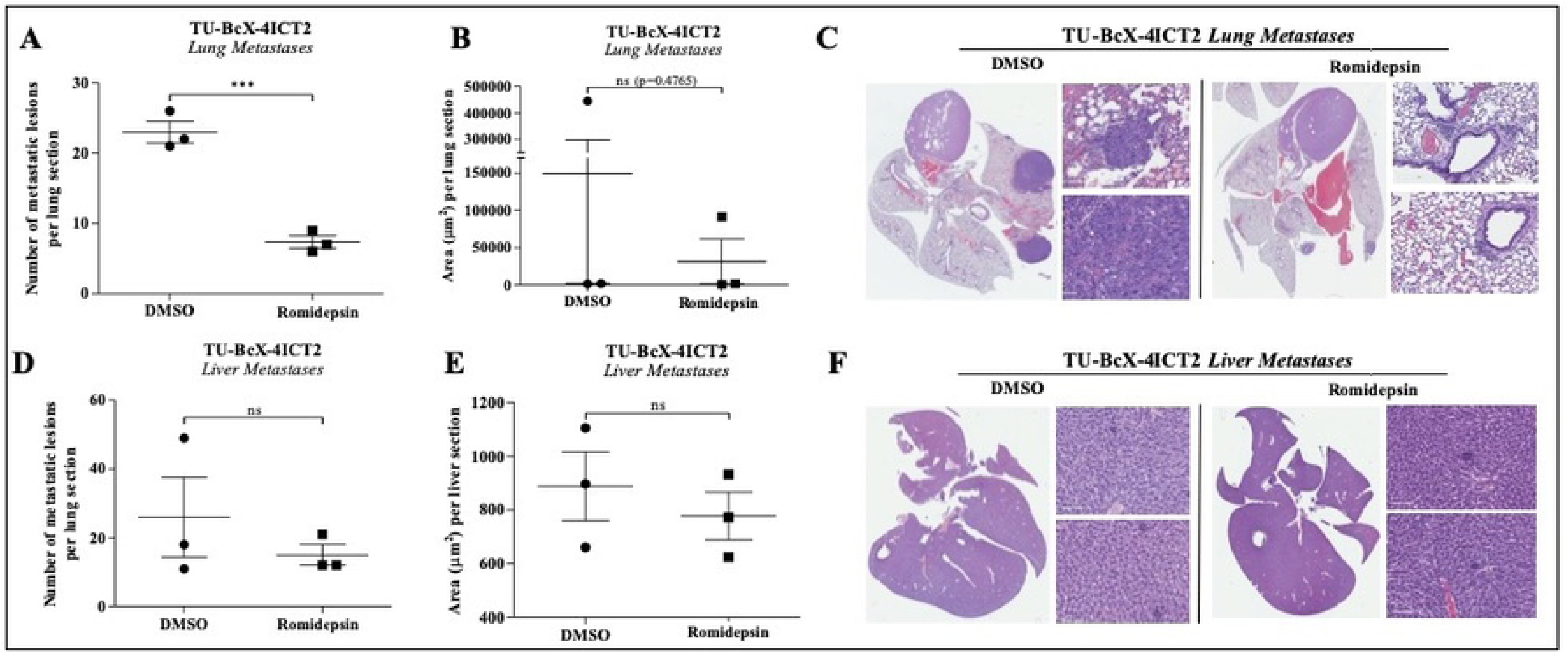
Romidepsin effect on metastasis in TU-BcX-4IC-implanted mice. Lungs and livers from mice implanted with TU-BcX-4IC and treated with romidepsin or DMSO vehicle were harvested, formalin-fixed, paraffin-embedded, and H&E-stained to visualize and quantify metastases. Romidepsin treatment (A) significantly decreased the number of metastatic lesions per lung section and (B) overall decreased the area of lung metastases compared to DMSO vehicle control. (C) Representative images of H&E-stained lungs in both treatment groups. (D) In liver sections, romidepsin (D) dramatically reduced the number of metastatic lesions per liver section and (E) overall decreased the area of liver metastases compared to DMSO. (F) Representative images of H&E-stained livers in both treatment groups. Error bars represent standard error of mean (S.E.M.). *** p<0.0001; ns = not significant.

### Romidepsin suppresses EMT-associated genes in MBC

RNA-sequencing of TU-BcX-4IC cells treated with romidepsin or DMSO demonstrated that romidepsin affected the expression of genes involved with EMT, extracellular matrix, and cell cycle signaling. In the EMT pathway, romidepsin interaction increased *CDH1*, *SNAI1* and *FOS* expression, and downregulated *PLAU*, *JDP2*, *TGFB11*, *WNT5B*, *NFKB1*, *WNT5A*, *FOSL1*, *WNT3*, and *ZEB2* in addition to other mesenchymal genes (Fig 4A, S2 Fig). In cell cycle signaling pathways, romidepsin increased *CDNK1A* expression and downregulated *HDAC7*, *PLK1*, *AURKB*, *PLK4*, *CENPA*, *AURKA*, *MKI67*, and *FOXM1* (Fig 4B). Romidepsin also affected ECM-related genes including upregulation of MMP9 and downregulation of *HAS2*, *PLAU*, *ADAMTS1*, *ECM2*, *ADAMTS5*, *DDR2*, *ADAM9*, *MMP14*, and *SERPINB1* (Fig 4C).

**Fig 4.**
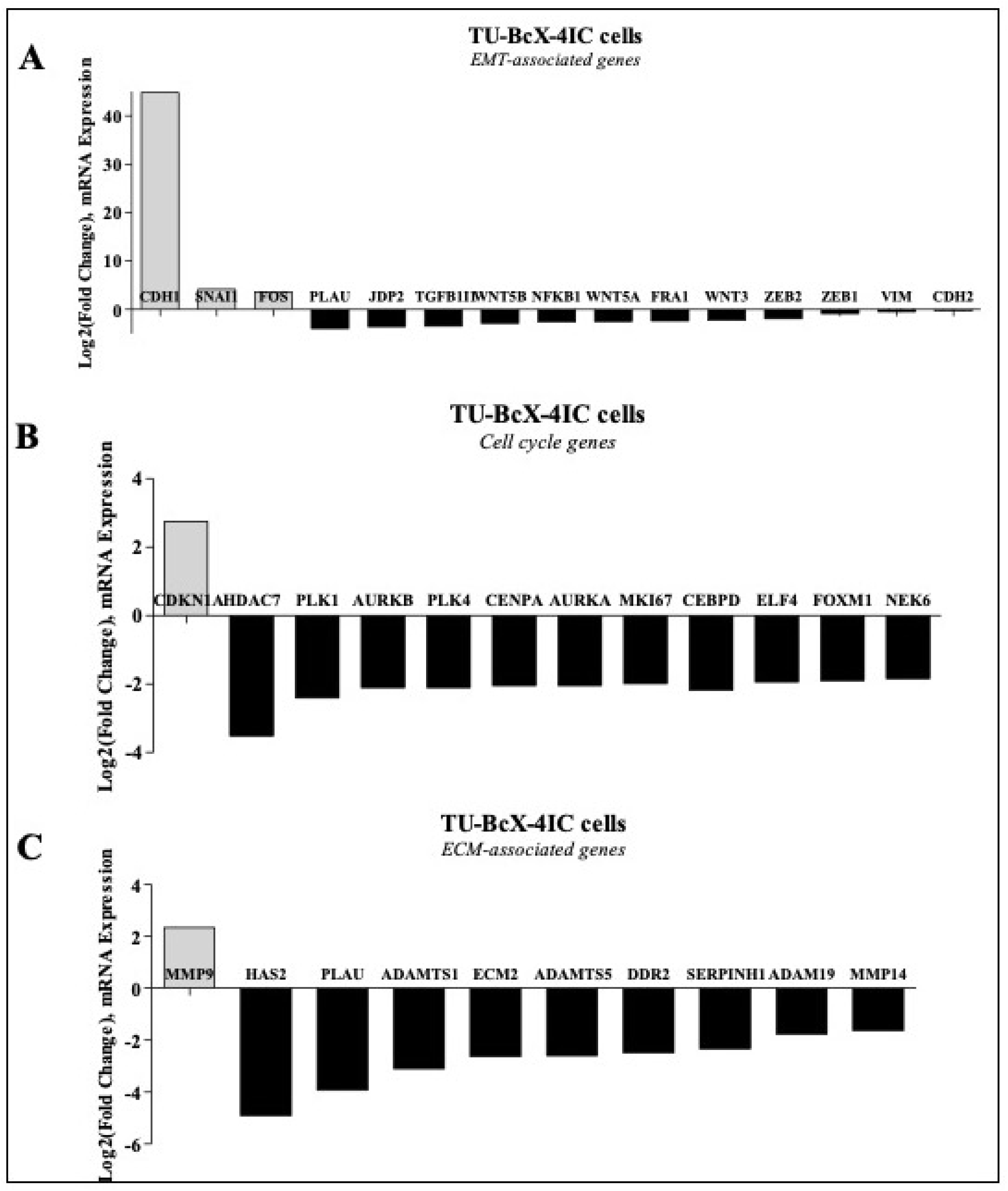
RNA sequencing analysis of romidepsin-treated TU-BcX-4IC cells compared to DMSO control. Romidepsin altered expressions of (A) EMT-related genes. Other cell signaling pathways that were affected include (B) cell cycle genes and (C) extracellular matrix-associated genes in TU-BcX-4IC cells. Data is shown as Log2 (fold change). mRNA expression of EMT genes (*CDH1*, *VIM*, *CDH2*, *ZEB1*, *ZEB2*) were analyzed using qRT-PCR.

We have previously shown that DACi suppresses the mesenchymal genes *CDH2*, *VIM*, *ZEB1*, and *ZEB2*, and increases the epithelial gene *CDH1* in TNBC cells (34-36). As mentioned previously, these genes were confirmed to be up- or downregulated in our RNA-sequencing analysis of TU-BcX-4IC cells treated with romidepsin compared to DMSO. We used qRT-PCR with romidepsin or DMSO to examine if similar gene changes were observed in other *in vitro* derivations of an MBC PDX model: primary cell lines, spheres, PDX-Os and *in vivo* tumor implants (Fig 5A-D, S3 Fig). Our data show romidepsin increases *CDH1* expression in the PDX-derived cell line, spheres, and PDX-Os. *CDH1* mRNA expression could not be observed in treated TU-BcX-4IC implanted tumors, due to low endogenous expression in the tumors. Across all studies, compared to DMSO, romidepsin reduced expression of the mesenchymal gene *VIM*, although to a lesser degree in PDX-Os. *CDH2* expression was downregulated by romidepsin in tumor implants but not in cells. *ZEB1* expression was not altered after romidepsin treatment in cells; *ZEB1* expression could not be observed in treated TU-BcX-4IC spheres, PDX-Os, nor tumor implants due to low endogenous transcript levels. Interestingly, *ZEB2* expression was downregulated in TU-BcX-4IC cells and spheres, although not in PDX-Os. This variability in gene expression suggests that other microenvironmental/stromal factors may play a role in the observed reversal of the mesenchymal phenotype caused by DACi.

**Fig 5.**
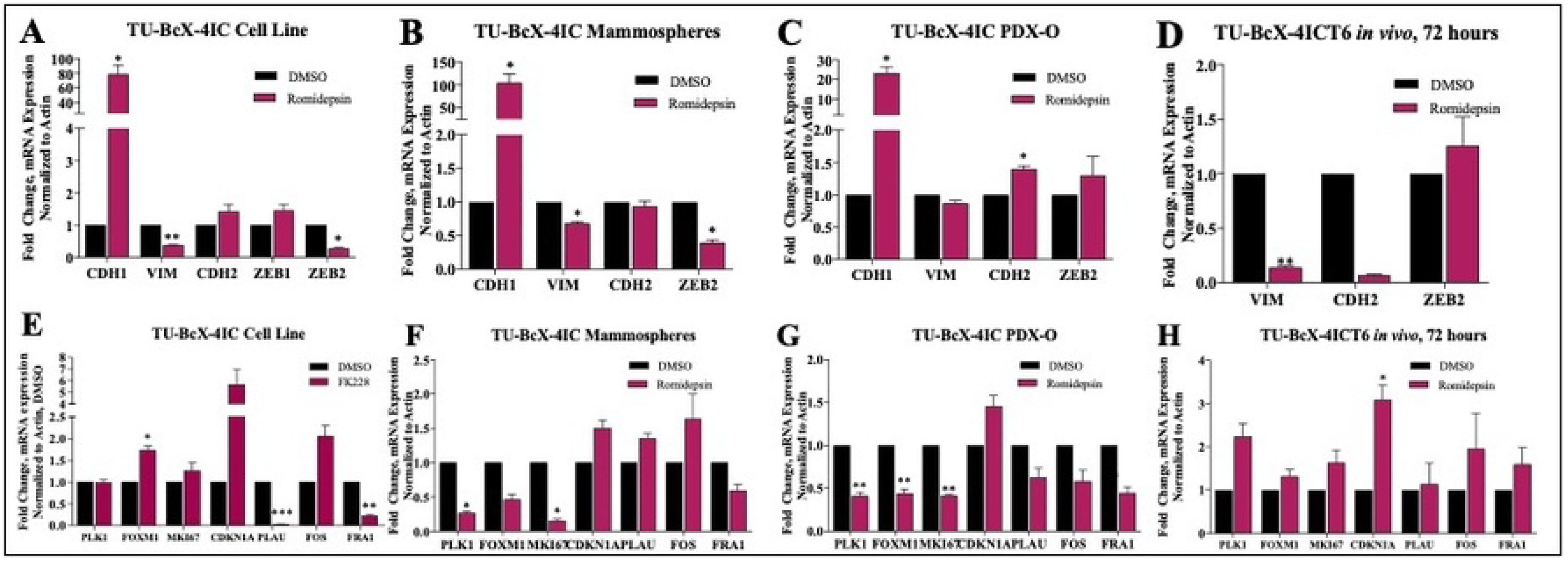
Effects of romidepsin on gene expression differs amongst treated cells, spheres, PDOs, and implanted tumors. Romidepsin treatment in TU-BcX-4IC (A) cell line, (B) spheres, (C) PDX-Os, and (D) *in vivo* tumor pieces implanted in SCID/Beige mice. qRT-PCR analysis was repeated with genes affected by romidepsin treatment compared to DMSO control based on RNA sequencing analyses (*PLK1*, *FOXM1*, *MKI67*, *CDKN1A*, *PLAU cFOS*, *FRA-1*). Again, romidepsin-treated TU-BcX-4IC (E) cells, (F) spheres, (G) PDX-Os, and (H) *in vivo* tumor pieces implanted in mice were used. Analyses were all normalized to β-actin and DMSO treated controls. Black bars represent DMSO; maroon bars represent romidepsin treatment (100nM, 72 hours).

Next, we examined changes in expression of additional EMT-, cell cycle-, and ECM-related genes that were significantly up- or downregulated in our sequencing analysis using the same derivations of the TU-BcX-4IC PDX model (Fig 5E-H, S3 Fig). Of the cell cycle genes assessed, *CDKN1A* was significantly upregulated in romidepsin-treated TU-BcX-4IC cells, spheres, PDX-Os and implanted tumors. The other cell cycle genes analyzed (*PLK1*, *FOXM1*, *MKI67*) were downregulated in romidepsin-treated spheres and PDX-Os but were not changed after romidepsin treatment of the cell line (*FOXM1* was increased in the cells) nor tumor implants. *PLAU* and *FRA1* were downregulated and while FOS was upregulated in the romidepsin-treated cells, consistent with the RNA-sequencing findings. However, *PLAU* expression was only also downregulated in PDX-Os, but not in implanted tumors nor in spheres. Conversely, *FOS* expression was also upregulated in romidepsin-treated spheres and implanted tumors, but not in PDX-Os (*FOS* has basically the same level of reduction than *PLAU* in PDX-O). *FRA-1* expression was downregulated in romidepsin-treated spheres and PDX-Os but had increased expression in implanted tumors. These findings show the differences in gene expression changes in cells compared to that in spheres, PDX-Os, and tumor implants; and thus, emphasize the importance of further analysis beyond the effects in monolayer cell populations in order to identify mechanisms of DACi on TNBC. In most derivations of our TU-BcX-4IC PDX model, romidepsin most consistently increased *CDH1* and *CDKN1A* gene expressions, and downregulated *VIM*, *ZEB2*, and *FRA1*.

### Differences in DACi effects on gene expression in PDX-Es derived from various passages in mice and temporal variability in in vivo treated tumor implants

PDX explants (PDX-Es) are translational models, because they maintain the tumor microenvironment and architecture that is unique to each patient tumor. Next, we tested drug effects on intact tumor PDX-Es. To examine how treatment of different passage tumor PDX-Es potentially affects our interpretations of DACi in MBC, we treated various passage PDX-Es with romidepsin (100 nM) for 72 hours. We tested the EMT-associated genes described previously (*CDH1*, *VIM*, *CDH2*, *ZEB2*), we found that romidepsin increased *CDH1* expression in PDX-Es, while *VIM* and *ZEB2* expression were not affected by romidepsin in any passage of PDX-Es. *CDH1* expression was downregulated in PDX-Es. With respect to the RNA-seq identified genes (*PLK1*, *FOXM1*, *MKI67*, *CDKN1A*, *PLAU*, *FOS*, *FRA1*), we observed that overall, the cell cycle genes *PLK1*, *FOXM1* and *MKI67* were consistently not altered after treatment of T2 nor T5 PDX-Es (there is increased expression of *FOXM1*). *CDKN1A* expression was upregulated PDX-Es, but not in T2 PDX-Es. *PLAU* expression was downregulated in T5 treated PDX-Es, but not in T2 (upregulated here). In T2 PDX-Es treated with romidepsin, *FOS1* was downregulated, but was upregulated in T5 treated PDX-Es. *FRA1* was downregulated in T2 and T5 PDX-Es. These data show inconsistencies with drug effects amongst treated intact tumor PDX-Es derived from various passages in mice.

Next, we sought to find short-term (72 hours) *in vivo* effects of romidepsin treatment on gene expression in the tumors compared to a longer-term (15 days) *in vivo* treatment. In both groups, suppression of *VIM* and *CDH2* (S4A-B Fig) and upregulation of *CDKN1A* mRNA expressions were observed, but these changes were most significant after short-term treatment (S4C-D Fig) suggesting changes in drug effectivity over time. Furthermore, *FRA1* was downregulated in both short- and long-term treatment groups, with a more significant downregulation in the long-term group (S4B Fig). There were also differences in gene expression in the short-term compared to long-term treatment groups. *FOS* gene expression was downregulated in tumors treated with romidepsin for 72 hours (S4C Fig), but not in those treated for 15 days (S4D Fig). Other genes (*ZEB2*, *FOXM1*, *PLAU*, *FOS*) were not affected after short-term nor long-term romidepsin treatments (S4A-D). Together, these data identify genes affected by short term treatment of DACi in MBC (*CDKN1A*, *VIM*, *CDH2*) and genes that were affected after prolonged treatment (*FOS*, *ZEB1*, *ZEB2*, *PLAU*) in TU-BcX-4IC tumors *in vivo*.

### Romidepsin pre-treatment sensitizes TU-BcX-4IC to NCI drugs

The link between EMT and drug sensitivity is an emerging area of research. Because we demonstrated that romidepsin suppresses the mesenchymal phenotype and thus, reverses EMT, we next evaluated if romidepsin sensitized TU-BcX-4IC cells to other targeted oncology agents. For these experiments, we used a set of clinically-approved oncology drugs provided by the National Cancer Institute (NCI). We found TU-BcX-4IC cells to be inherently resistant to many of the drugs provided in the panel (Fig 6A). However, in an initial screen where we pre-treated TU-BcX-4IC cells with romidepsin (50 nM) resulted in markedly increased sensitivity to select drugs (Fig 6A). Pre-treatment with romidepsin increased sensitivity to HER-targeted inhibitors (afatinib, lapatinib), the estrogen-receptor inhibitor fulvestrant, and MEK1/2 inhibitors (trametinib, cobimetinib). Furthermore, pre-treatment increased sensitivity to other targeted inhibitors dabrafenib (BRAF), everolimus (mTOR), ponatinib (Bcr/Abl), crizotinib (c-MET, RON, ALK) and bleomycin (Fig 6A). Together, these data show that DACi adjuvant treatment sensitizes drug-resistant TU-BcX-4IC cells to clinically approved drugs. We also show drugs that were cytotoxic to both pre-treatment naive and romidepsin pre-treated cells, including bortezomib, mitoxantrone, topotecan, and epirubicin (S5 Fig). Then, we selected the two MEK1/2 inhibitors, cobimetinib and trametinib, that were sensitized with romidepsin-treated cells to further interrogate the effects of adjuvant romidepsin treatment. To confirm our findings that romidepsin sensitizes TNBC cells to MEK1/2 targeted therapy, we first replicated the initial screen by pre-treating TU-BcX-4IC cells with romidepsin for 24 hours before treating cells with cobimetinib and trametinib. TU-BcX-4IC cells pre-treated with romidepsin for 24 hours had increased sensitivity to the MEK1/2 inhibitors compared to romidepsin or MEK1/2 inhibition alone (Figs 6B,6D). Co-treatment (cells treated at the same time point) with romidepsin and cobimetinib or trametinib amplified a similar mesenchymal-to-epithelial transition phenotypic response that we observed in our initial morphology experiments (Fig 6C, S6 Fig). The TU-BcX-4IC cells appeared more epithelial morphologically: round-shaped cells with higher cytopSlasm-to-nuclear ratios.

**Fig 6.**
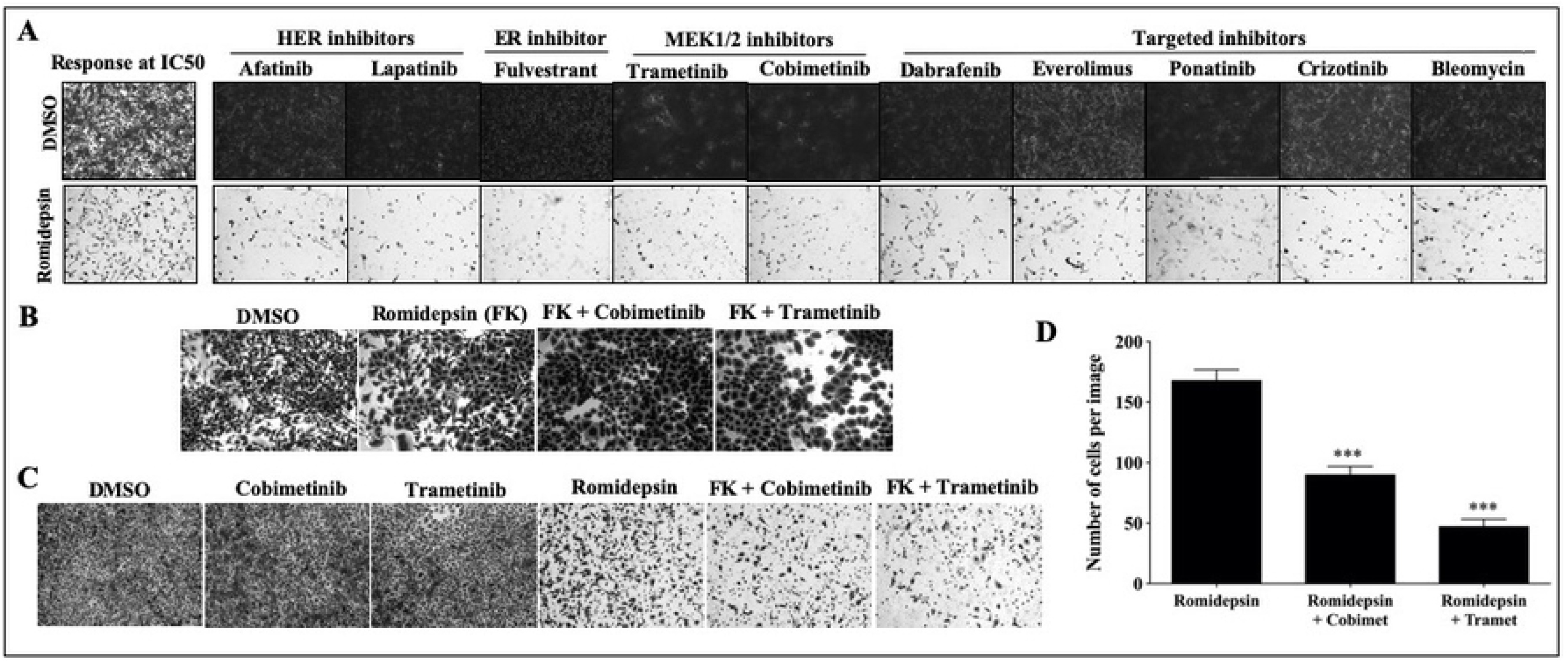
Romidepsin sensitizes TU-BcX-4IC cells to select oncology drugs. TU-BcX-4IC cells were treated only with agents from the NCI-approved oncology set or pre-treated with romidepsin (50 nM, 72 hours) prior to the oncology agents. Crystal violet staining was performed to visualize differences in cell viability between the groups. (A) Romidepsin sensitized TU-BcX-4IC cells to HER-targeted inhibitors (afatinib, lapatinib), the ER inhibitor fulvestrant, MEK1/2 inhibitors (trametinib, cobimetinib), as well as other targeted inhibitors (dabrafenib, everolimus, ponatinib, crizotinib), and bleomycin. Representative images are shown at 40X magnification. Response of both treatment groups to fluorouracil is shown as an example of baseline response to romidepsin, as well an example of 50% response of TU-BcX-4IC cells. (B) TU-BcX-4IC cells were pre-treated with romidepsin (50 nM) for 24 hours before treatment with cobimetinib and trametinib (1 µM) for an additional 24 hours. (C) TU-BcX-4IC cells were co-treated with romidepsin (50 nM) and cobimetinib or trametinib (1 µM) for 72 hours. (D) Quantification of cells remaining when cells were treated with romidepsin alone or pre-treated with romidepsin for 24 hours followed by cobimetinib or trametinib. Notably, the abundance of cells in the DMSO group prevented quantification of this treatment control and is not included in the graph. ***p < 0.001.

## Discussion

Metaplastic breast cancer is a rare breast cancer subtype, for which there are limited therapeutic options. Clinically, MBCs exhibit rapid tumor growth rates and are chemo-refractory to a variety of chemotherapies [60]. There is a limited understanding of the underlying pathology and biological pathways that drive this proliferative tumor type, due to the rarity of MBC. Given the current treatment options with limited efficacy and poor response rates, insights into new therapeutic strategies or regimens for MBC are crucial. EMT is a proposed mechanism for the initiation of metastasis and acquisition of drug resistance in TNBC and MBC; pharmacologic inhibition of this transition is one proposed mechanism to prevent metastasis and resistance. We previously published that DACi reverses the mesenchymal morphology and mesenchymal gene expressions in aggressive TNBC subtypes [35, 36]. In this study, the response of MBC to DACi treatment was evaluated.

TU-BcX-4IC tumors implanted *in vivo* and TU-BcX-4IC patient-derived cells *in vitro* exhibit highly tumorigenic and metastatic capacities that model the clinically aggressive nature of MBC. Thus, this new PDX model is an ideal tool for studying drug effects on the dynamic processes that occur in resistant cancers. While DACi treatment in our MBC model suppressed some aspects of a mesenchymal cell phenotype, we did not observe the canonical EMT phenotypic change that we had previously observed with DACi treatment in other TNBC cells. Both inhibitors were cytotoxic *in vitro* to TU-BcX-4IC cells and PDX-Os, and IC-50 concentrations differed with the DACis. Given the potential undesired side effects of using pan-DACi agents, in this study, we chose to focus on the response of TU-BcX-4IC to romidepsin, the targeted HDAC1/2 inhibitor. In previous studies, we found that romidepsin suppressed tumor growth and metastasis of established TNBC cell lines and implanted TNBC PDX tumors *in vivo* [34–36]. Romidepsin treatment did not inhibit growth of TU-BcX-4IC implanted tumors in murine models, nor did treatment affect average area of lung metastases nor number and area of liver metastases compared to DMSO control-treated tumors. Notably, romidepsin treatment significantly reduced number of lung metastases compared to vehicle control, suggesting romidepsin prevents direct metastatic seeding to the lungs. The presence of cells that break free from the primary tumor and circulate before seeding in distal tissue sites has been investigated as a predictive measure for metastasis [62, 63]. It was also observed that romidepsin suppressed the number of circulating tumor cells (CTC) in the peripheral blood of mice implanted with TU-BcX-4IC tumors compared to vehicle control. Romidepsin treatment suppressed both MBC cell migration and mesenchymal gene expression. Together, these data support our working hypothesis that HDAC1/2 promotes lung tropism through regulation of the cells’ capabilities to escape the primary tumor site and extravasate to the lungs. A limitation to our study was that one model was used; due to the rarity of this malignancy, testing DACi in other MBC PDX models are necessary to ascertain broader understanding of a role for DACi in MBC tumor growth and metastasis.

This project was initiated to evaluate DACi therapy in MBC. Throughout the course of our experiments, we employed various derivations of PDX models to interrogate this objective. Profound observations were noted, which led us to pursue a secondary focus of this project: examining drug effects on different variations of PDX models. Depending on the cell or tumor system used, DACi had different effects on gene expressions. This resulted in contradictory conclusions pertaining to how DACi affected EMT and cell cycle genes in TNBC and MBC. These findings emphasized the importance of integrating multiple cell and tumor systems in drug discovery research, in order to draw more accurate conclusions regarding a drug’s role in cancer. We compared DACi treatment in the patient-derived cell line, cells grown in suspension, PDX-Os, tumor pieces, and implanted intact tumors *in vivo*. We examined both EMT-related genes *(CDH1, VIM, ZEB1, ZEB2, CDH2, PLAU, FOS, FRA1)* [34, 35] and, since DACi regulates cell proliferation, cell cycle-related genes (*PLK1*, *FOXM1*, *CDKN1A, MKI67*). Patterns were observed in genes that were consistently affected by all patient-derived models, including *CDH1, VIM, FRA1* and *CDKN1A.* Other genes were only affected in some models, including *ZEB2* which was downregulated in the cell line and spheres. These data demonstrate response to romidepsin varied depending on PDX-derived model used.

An interesting observation was that overall, gene expressions affected by romidepsin treatment in the 3D cultured spheres derived from the TU-BcX-4IC cell line were most similar to that of PDX-Os. Also, in the more translational models (PDX-Os, PDX-Es, and implanted tumors), there was reduced expression of *PLAU, FOS* and *FRA1* in the implanted tumors compared to the PDX-Os and PDX-Es. Our findings are consistent with other groups that have found drug sensitivity and associated gene expression changes differ in adherent compared to suspension culture systems, and *in vivo* analyses [55,56,64,65]. One limitation in therapeutic discovery research is that laboratory findings are not often translated into clinical practice [61], and we believe integrating multiple derivations of PDX models is a step in overcoming this limitation.

We recognized difficulty in comparing drug effects on gene changes in tumors treated *in vivo*, which often occur in long time increments (15 days), to cells and tumor explants treated *in vivo* and *ex vivo* in shorter time increments (72 hours). To address this and to evaluate the temporal relationship between DACi treatment and its effects, we treated mice implanted with TU-BcX-4IC tumors for 72 hours *in vivo* and compared these findings to the long-term treatment group. While some genes remained unchanged, there were important differences, which supports our hypothesis that romidepsin has both immediate and later-onset effects to suppress metastasis and alter downstream signaling pathways. Unique genes that were affected by short-term compared to longer-term treatments include upregulation of *CDKN1A* and suppression of *VIM* and *CDH2* expression. After long-term treatment, downregulation of genes for transcription factors (*FOS*, *ZEB1*, and *ZEB2)* and genes encoding proteins involved in cytoskeletal aspects of the mesenchymal morphology in EMT (*VIM* and *CDH2)* was observed. Because *VIM* and *CDH2* were more immediately downregulated compared to the transcription factors, combined with our findings that romidepsin promotes epithelial phenotypes after 72 hours, it can be inferred that romidepsin has an early effect on reversal of mesenchymal morphology. Then, romidepsin further suppresses metastatic effects after prolonged treatment by suppressing the transcription factors that drive the mesenchymal phenotype.

EMT has emerged as a potential mechanism of developing drug resistance in cancer [41, 46]; this is especially important in MBC due to de novo resistance to multiple cytotoxic therapies. Given our findings that DACi suppressed the mesenchymal phenotype, we investigated the role of DACi on chemoresistance using an NCI oncology drug set. TU-BcX-4IC cells were sensitized to various anti-cancer drugs including bleomycin, HER inhibitors afatinib and lapatinib, ER inhibitor fulvestrant, MEK1/2 inhibitors trametinib and cobimetinib, and other targeted inhibitors (dabrafenib, everolimus, ponatinib, and crizotinib) following pre-treatment with romidepsin. These data suggest that histone DACi increases TU-BcX-4IC susceptibility to some oncology drugs. In addition, romidepsin improved TU-BcX-4IC cell sensitivity to bortezomib, mitoxantrone, topotecan, and epirubicin. The reversal of drug resistance after pre-treatment with romidepsin suggests preliminary evidence that DACis serve as additional therapeutic agents to current anti-tumorigenic drugs to maximize the therapeutic potential in highly resistant cancers. Two compounds were selected from the NCI drug panel set (cobimetinib and trametinib, MEK1/2 inhibitors) that, when combined with romidepsin, elicited a cytotoxic response to TU-BcX-4IC cells. These data implicate a synergistic role of DACi treatment with MEK1/2 inhibitors in MBC. This is consistent with previously published data regarding a synergistic role of DACi and MEK1/2 therapy in various tumor types. Some examples include the combined effects that induce fatality of leukemia cells [66], combined therapy resulted in attenuated tumor growth of BRAF-mutated colorectal cells [67], and DACi treatment enhanced the antitumor effects of MEK1/2 treatment in RAS-mutated lung cells [68].

In this study, we sought to evaluate DACi therapy in MBC using a new PDX model established by our group. When multiple derivations tumors and cells derived from the same PDX model were employed in mechanistic studies, the focus of the investigation shifted to emphasize the importance of using various cell and tumor systems to acquire a more complete and translational understanding of targeted drug response in complex breast cancer subtypes. Our findings are important in preclinical studies that evaluate the effects of novel therapeutic agents in all cancers and are not restricted to breast cancer. The application of multiple cancer models is necessary to more accurately translate preclinical observations into clinical practice.

## Conclusion

The results of this study showed DACi suppression of migration and mesenchymal phenotypes in MBC, compared to DMSO controls. We demonstrated DACi regulation of the epithelial-mesenchymal transition axis and suppression of the CTC population. Our results serve as preliminary evidence that DACi treatment in combination with MEK1/2 inhibitors exerts a synergistic effect in MBC. Our results suggest that cancer models can serve as an important preclinical tool to identify effective therapeutic agents for complex breast cancer subtypes.

## Acknowledgements

Our group is very appreciative of the Genomics Core, under the direction of Dr. Erik Flemington, at Louisiana State University Health Sciences Center for performing the RNA sequencing. We also appreciate the collaboration with Alan Tucker in the Flow Cytometry and Cell Sorting Core Lab of Tulane Center for Stem Cell Research and Regenerative Medicine. We would also like to thank the Histology Core facilities at Tulane University for the hematoxylin and eosin stains of the tissue from the mice experiments. We would also like to thank the animal vivarium staff at Tulane University for their advice and assistance with the mice experiments. We are always grateful for the support, both financially and socially, of the Krewe de Pink organization in New Orleans. Finally, but most importantly, we are thankful to patients who donated their breast cancer tissue specimens to benefit breast cancer research; their contributions are valued and appreciated.

## Supporting Information

**S1 Fig.**
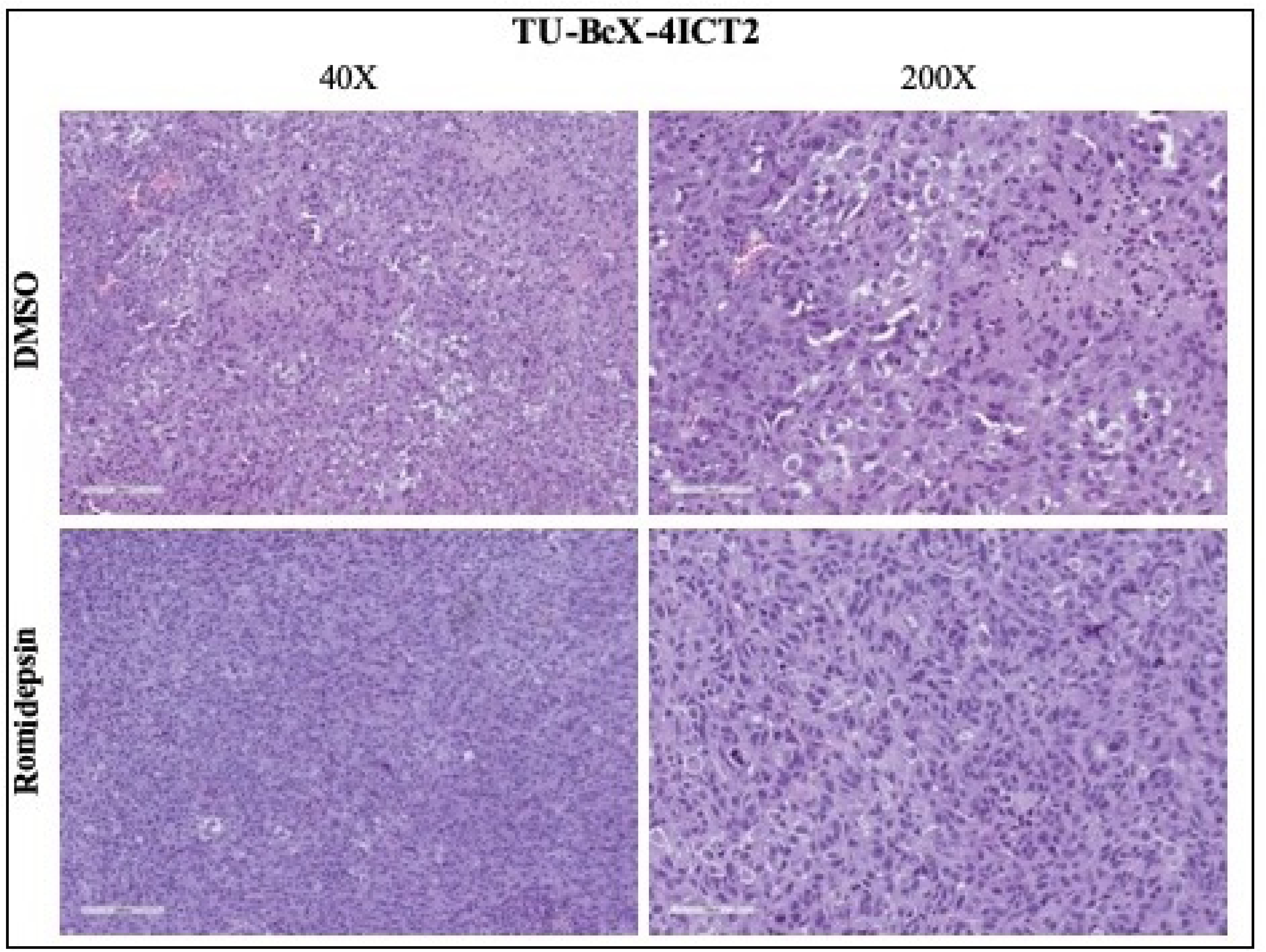
Romidepsin cytotoxicity to tumors in patient-derived xenograft model. Tumors were excised, formalin-fixed, paraffin-embedded and H & E-stained to visualize cellular composition and changes in romidepsin-treated tumors compared to DMSO control. Atypical histologic features were seen on both romidepsin and DMSO tumor specimens, including mitotic figures, nuclear pleomorphism, and hyperchromicity.

**S2 Fig. Romidepsin modifies networks of EMT-related genes in TU-BcX-4IC cells compared to DMSO.**

Pathway analyses demonstrate the networks of EMT-related gene changes in romidepsin-treated TU-BcX-4IC cells compared to DMSO control treated cells. Data is shown as Log2 (fold change). Genes highlighted by green represent upregulated genes and downregulated genes are highlighted in red.

**S3 Fig. Romidepsin suppresses expression of EMT-associated genes and gene expression differs amongst treated cells, mammospheres, PDX-Os, PDX-Es, and implanted tumors.**

All data is shown as fold change ± SEM normalized to DMSO treatment controls.

**S4 Fig. Effect of short-term treatment with romidepsin compared to long term effects on gene expression.**

Expression of EMT mRNAs (CDH1, VIM, CDH2, ZEB1, ZEB2) were analyzed using qRT-PCR. Romidepsin was treated in TU-BcX-4IC tumor pieces implanted in SCID/Beige mice. Drug effect studies were (A) short term (72 hours) or (B) long term (15 days). qRT-PCR analysis was repeated with genes affected by romidepsin treatment compared to DMSO control based on RNA sequencing analyses (PLK1, FOXM1, MKI67, CDKN1A, PLAU, cFOS, FRA-1). Drug effect studies were (C) short term (72 hours) or (D) long term. Black bars represent DMSO; maroon bars represent romidepsin treatment (100 nM, 72 hours). All experiments were run in triplicate. Error bars are shown as S.E.M.

**S5 Fig. TU-BcX-4IC cells were treated with oncology drugs selected from the NCI drug set cytotoxic to TU-BcX-4IC cells and compared to cells pre-treated with romidepsin.**

Crystal violet staining of TU-BcX-4IC cells pre-treated with romidepsin for 48 hours (50 nM), or without romidepsin, and then subsequently treated with the NCI oncology drug set. “FK” denotes FK228, or romidepsin, treatment.

**S6 Fig. Quantification of co-treatment studies with romidepsin and MEK 1/2 inhibitors (cobimetinib or trametinib) and pre-treatment studies with romidepsin.**

TU-BcX-4IC cells were concomitantly treated with romidepsin and a MEK 1/2 inhibitor. Pre-treatment with romidepsin was studied in TU-BcX-4IC cells by either pre-treating with romidepsin (50 nM) or not pre-treating for 48 hours, and then treating with cobimetinib or trametinib (1 µM). Crystal violet stained cells were lysed and absorbance was measured at 570 nm to quantify staining results.

